# The under-recognized dominance of magnetosome gene cluster-containing bacteria in oxygen-stratified freshwater ecosystems

**DOI:** 10.1101/2024.04.27.591411

**Authors:** Runjia Ji, Juan Wan, Pranami Goswami, Jianxun Shen, Yonxin Pan, Wei Lin

## Abstract

Magnetotactic bacteria (MTB) capable of magnetosome organelle biomineralization and magnetotaxis are widespread in chemically stratified aquatic environments. Conventionally, it has long been considered that the overall abundance of MTB in microbiota is not very high and that Magnetococcia is the most frequently identified and predominant MTB members. However, the diversity and distribution of MTB in chemically stratified environments remain elusive due to the lack of large-scale systematic analyses. Here we conduct a comprehensive survey of genomes containing magnetosome gene clusters (MGCs), a group of genes responsible for magnetosome biomineralization and magnetotaxis, in 267 metagenomes from 38 oxygen-stratified freshwater environments. A total of 63 MGC-containing genomes belonging to eight bacterial phyla are reconstructed, including the newly identified Myxococcota. We discover an unexpectedly high relative abundance of putative MTB (up to 15.4% of metagenomic reads) in hypoxic and anoxic water columns, in which Deltaproteobacteria, rather than traditionally considered Magnetococcia, are the most ubiquitous and predominant MGC-containing bacteria. Our analysis reveals a depth-specific taxonomy and function of MGC-containing bacteria in stratified water columns shaped by physicochemical conditions. These findings underscore the unrecognized ecophysiological importance of MTB in freshwater ecosystems.

## Introduction

Diverse organisms can sense Earth’s magnetic field for navigation and orientation^1,2^. Magnetotactic bacteria (MTB) are so far the only known group of prokaryotes with the ability of magnetoreception^3–5^. Magnetoreception in MTB (also known as magnetotaxis) is triggered by magnetosomes, a unique bacterial organelle that consists of membrane-bounded nano-sized magnetite (Fe_3_O_4_) and/or greigite (Fe_3_S_4_) crystals. The arrangement of magnetosomes in chain-like structures optimizes the magnetic dipole moment of MTB cells, facilitating their passive alignment with the geomagnetic field. The biosynthesis of magnetosomes is regulated by a collection of magnetosome genes^6^, which are in close proximity within MTB genomes, called magnetosome gene clusters (MGCs)^7^. Magnetosomes are so far the only confirmed magnetoreceptors, making MTB an accessible and relatively simple model system for studying the origin and evolution of magnetoreception and biomineralization. Moreover, magnetosomes as magnetic nanoparticles with unique characteristics and high biocompatibility have great potential for bio– and nanotechnology applications^8–11^.

Stratification is an important physicochemical structuring of aquatic environments, of which oxygen-stratified environments are one of crucial habitats for understanding climate change and its impact on ecosystems^12,13^. These environments not only have an important impact on the diversity and ecology of microbial communities, but also represent hotspots for discoveries of novel microorganisms^14^. MTB are a typical group of microorganisms that mainly thrive in proximity to the oxygen-stratified aquatic environments, that is, the oxic-anoxic transition zone (OATZ) ^3,15–17^. Previous studies reported that Fe_3_O_4_-producing MTB are normally present in hypoxic to anoxic layers in the OATZ, while Fe_3_S_4_-producing MTB generally inhabit the anoxic sulfidic regions below the OATZ^18–21^. MTB are considered to be non-negligible ecosystem engineers in stratified aquatic habitats due to their worldwide distribution and important roles in the global cycling of iron and several other elements^22,23^. However, mechanisms underneath vertical distributions of various MTB groups and their functional profiles in stratified habitats remained largely uncharted due to the lack of extensive (meta)genomic analysis.

The diversity and phylogeny of MTB have recently been greatly expanded and, to date, MTB are found in at least 16 bacterial phyla^23–25^. The ongoing discovery of new MTB lineages suggests that our knowledge of their diversity and environmental function remains very limited. The classic method for studies of MTB diversity and ecology takes advantage of their magnetotactic behavior and pre-processes by magnetic enrichment of MTB cells from environmental samples prior to further analysis. Through this method, Magnetococcia within the phylum Proteobacteria (recently renamed Pseudomonadota) have long been viewed as the predominant MTB members in most freshwater and marine environments^26–31^. While members of the Nitrospirota are occasionally the dominant MTB populations in some ecosystems^24,32^. However, the magnetic enrichment process depends on magnetotaxis capability and motility of cells, possibly resulting in a biased understanding of MTB diversity and distribution^33^.

Here, to better profile the taxonomic and functional diversity of MTB in oxygen-stratified water columns, we conducted a genome-resolved metagenomic analysis of 267 freshwater metagenomes without magnetic enrichment pre-processing across the Northern Hemisphere. To our knowledge, this is the first comprehensive metagenomic survey of MGC-containing bacteria from a series of chemically stratified water columns, which revealed that MGC-containing bacteria occupy up to 15.4% of local microbiotas in hypoxic and anoxic water columns, indicating a largely underrated ecological role of putative MTB. Deltaproteobacteria are of strikingly high ubiquitous and predominant MGC-containing groups across these samples. In addition, the vertical distribution along with the metabolic function prediction of dominant MGC-containing members across stratified water columns were investigated, uncovering mechanisms that shaped the vertical niche differentiation pattern of putative MTB communities.

## Results

### Metagenomic dataset description

To comprehensively investigate the putative MTB population in freshwater environments, we used a publicly available metagenomic dataset (Project accession: PRJEB38681) that was reported and deposited to the European Nucleotide Archive (ENA) by Buck et al^34^. A total of 267 metagenomes from 38 oxygen-stratified freshwater environments (including 25 lakes, 12 ponds, and one reservoir) were analyzed in this study (Fig. 1a and Supplementary Table S1). These samples were collected from five countries and regions across tropical to polar zones of the Northern Hemisphere (Fig. 1b). 225, 34, and 8 of these metagenomes were collected from the freshwater lakes, ponds, and reservoir, respectively (Fig. 1c and Supplementary Table S1). Except for 9 metagenomes from Lake Alinen Mustajärvi sediments, all other metagenomes were collected from water columns (Fig. 1c and Supplementary Table S1). In total, the dataset encompassed over 9.5 billion paired-end Illumina sequencing reads, corresponding to a total of 2.8 terabases (150 bp/reads × 9.5 × 2 billion reads).

**Fig. 1.**
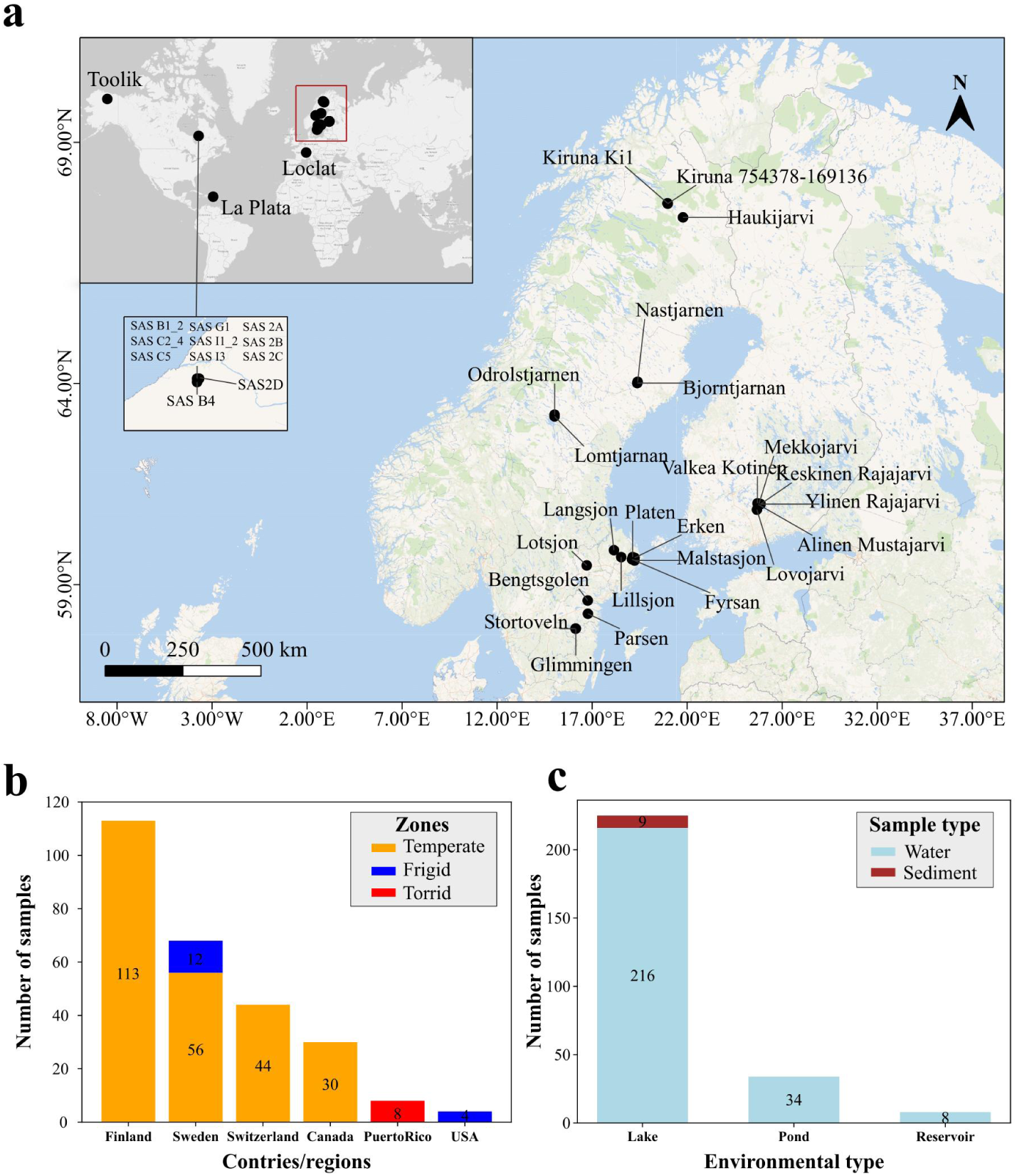
Distribution and overview of sampling sites. **a** Global distribution of 38 oxygen-stratified lakes and ponds selected in this study^34^. The base map depicts the region indicated by the red squared region in the insert. 11 lakes and ponds with close latitude and longitude in Canada were shown in the small inserted map. **b** Geographic distribution of 267 metagenomes analyzed in this study. Samples were collected from six countries/regions from the frigid zone to the tropical zone, with the largest number of samples collected from the temperate zone (Supplementary Table S1). **c** Environmental types of 267 metagenomes. Most samples were collected from water bodies of lakes, ponds, and one reservoir, and only nine were sediment samples from Lake Alinen Mustajärvi (Supplementary Table S1).

### Expansion of magnetotaxis potential in the Bacteria domain

After sequence assembly, binning, and MGC screening (see Methods for details), a total of 63 MGC-containing genomes were obtained. Among them, 27 genomes were identified as high quality (completeness ≥90% and contamination <5%) and 36 as medium quality (completeness ≥50% and contamination <10%) according to the minimum information about a metagenome-assembled genome (MIMAG) standard^35^ (Supplementary Table S2). Using an average nucleotide identity (ANI) cutoff of 99%^36^, 63 MGC-containing genomes were dereplicated into 41 non-redundant (nr) genomes, of which 16 were high-quality and 25 were medium-quality (Supplementary Table S2). Based on the GTDB release R207, the 41 nr MGC-containing genomes are classified into eight bacterial phyla: Desulfobacterota (16 genomes), Pseudomonadota (12 genomes), Omnitrophota (5 genomes), Bdellovibrionota (4 genomes), Desulfobacterota_I (1 genome), Desulfobacterota_G (1 genome), Myxococcota (1 genome), and Hydrogenedentota (1 genome) (Fig. 2, Supplementary Fig. S1, and Supplementary Table S2). One MGC-containing genome belonging to the phylum Myxococcota is reported here for the first time (SFLP_MTB_KT4_bin17), suggesting a potentially much more widespread distribution of MTB in the Bacteria domain. Besides, 15 of the 41 nr MGC-containing genomes could not be assigned at the species level, and two could not be assigned at the genus level (Supplementary Table S2), indicating that many of these genomes represent novel populations. Contrary to previous thought that Magnetococcia are often the dominant MTB members, here only one genome (SFLP_MTB_Umea3p4_bin52) belonging to Magnetococcia was detected (Fig. 2 and Supplementary Table S2).

**Fig. 2.**
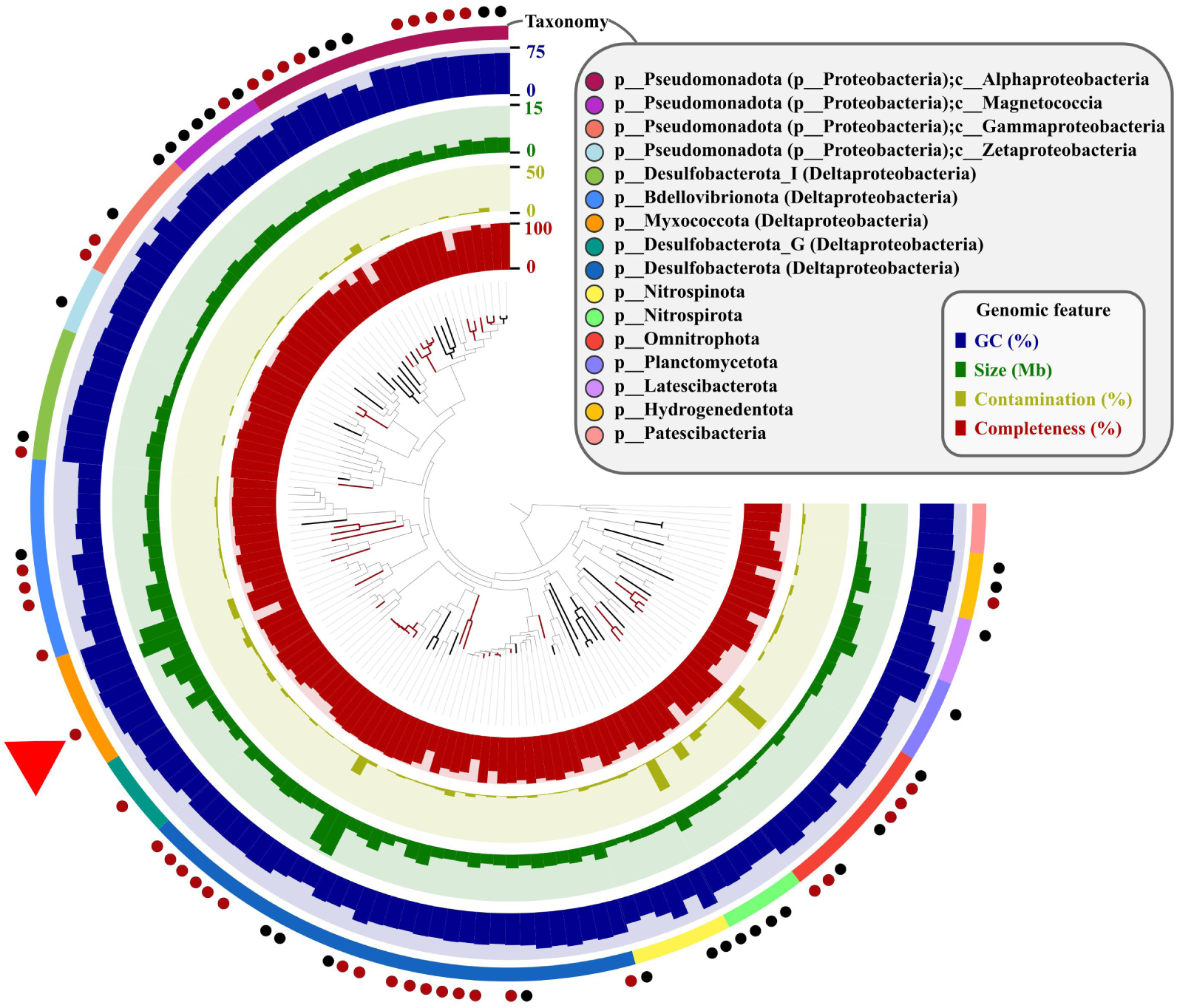
Phylogeny and quality of reconstructed MGC-containing genomes. Phylogenomic tree of 41 non-redundant genomes reconstructed here together with 33 previously published representative MTB genomes and 60 non-MTB genomes constructed using 120 bacterial single-copy marker protein sequences. From innermost to outermost, layers surrounding the phylogenomic tree indicate the completeness, contamination, size, GC content, and taxonomy at the phylum and class level of each genome, respectively. The branches of 41 non-redundant MGC-containing genomes obtained here are colored in red and signified by red dots on the outer edge. The branches of 33 previously published representative MTB genomes are colored in black and signified by black dots on the outer edge. The MGC-containing genome affiliated to the phylum Myxococcota is indicated with a red triangle.

All 41 nr MGC-containing genomes reconstructed in this study contain Fe_3_O_4_-type MGCs and their magnetosome gene composition and arrangement showed a distinct lineage-specific feature. The magnetosome gene content and organization are in general more conserved within phylogenetically close members than those of distantly related lineages (Fig. 3). For example, the overall structures of MGCs within the Deltaproteobacteria (recently reclassified into four phyla: Desulfobacterota, Bdellovibrionota, Myxococcota, and SAR324^37^) are similar. However, we noted that although Myxococcota is phylogenomically close to Bdellovibrionota (Fig. 2, Supplementary Fig. S1), the MGC of the Myxococcota genome (SFLP_MTB_KT4_bin17) has a unique gene order that differs from those within Bdellovibrionota and other phyla (Fig. 3).

**Fig. 3.**
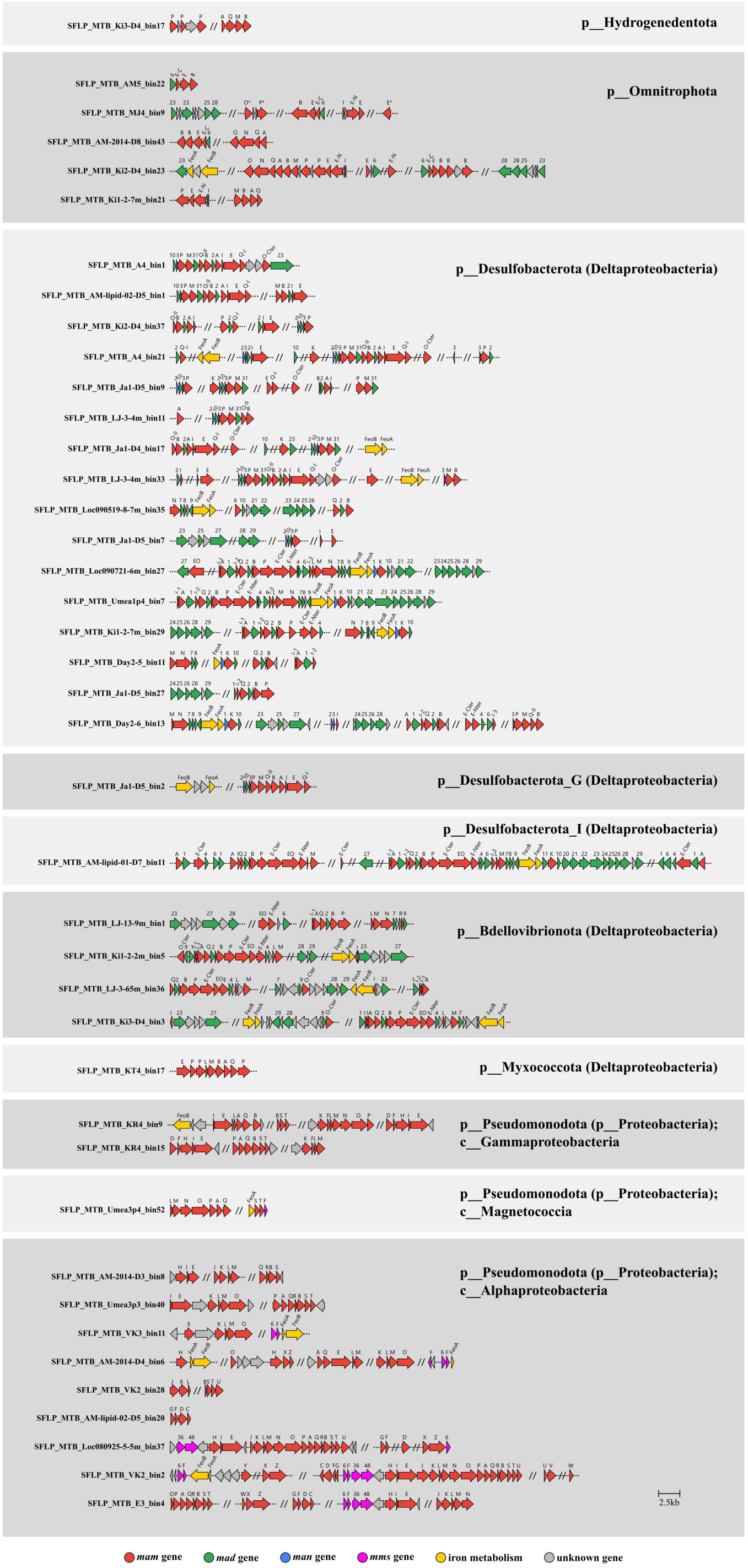
Magnetosome gene clusters (MGCs) of 41 non-redundant MGC-containing genomes obtained in this study. The names are indicated on the left and different lineages are on the right. All identified MGCs in this study are Fe_3_O_4_-type.

### Unexpectedly high relative abundance of MGC-containing bacteria in freshwater microbiota

To assess the distribution and relative abundance of putative MTB communities in freshwater ecosystems, we performed genome-wide quantitative read recruitment analysis using the 41 nr MGC-containing genomes across all metagenomes (see Methods for details). Based on a detection threshold of 0.25^38^, putative MTB were identified in 142 metagenomes obtained from 22 lakes and ponds in Canada, Finland, Switzerland, and Sweden (Supplementary Table S3). The remaining 16 lakes and ponds, where no MGC-containing genome was detected based on the analysis here, primarily contain surface samples characterized by relatively high levels of dissolved oxygen. Therefore, while no MGC-containing genome detected in these 16 lakes and ponds, it is likely that the absencis might attributed to inadequate sampling of low-oxygen aera, thus the presence of MTB in those environments cannot be entirely ruled out.

The relative abundance of MGC-containing bacteria across lakes and ponds was then profiled. Samples collected on the same day from the same lake/pond were grouped together (referred to as lake-time groups). In this way, 267 samples were categorized into 80 lake-time groups (Supplementary Table S4). The number of total samples (N_t_) and the number of putative MTB-containing samples (N_mtb_) of each lake-time group were counted. After discarding the lake-time groups with N_t_ < 3 or N_mtb_ < 2 (denoted as inadequate sampling groups), 26 lake-time groups containing 165 samples passed the filtering (Fig. 4 and Supplementary Table S4). For each metagenome, the percentage of reads retrieved during read recruitment was used as a proxy for the relative abundance of MGC-containing bacteria. Within each lake-time group, the sample with the highest relative abundance was considered the optimal niche for putative MTB community, and the percentage value was used to represent the relative abundance of MGC-containing bacteria in the microbiota of that group. Our results showed that the relative abundance of MGC-containing bacteria ranged from 0.5% to 15.4%, with a median of 3.5% across the 26 lake-time groups (Fig. 4 and Supplementary Table S4). These results highlight the substantial abundance of MGC-containing bacteria in freshwater ecosystems, indicating a much more important ecological role of putative MTB in biogeochemical cycling of freshwater ecosystems than previously thought.

**Fig. 4.**
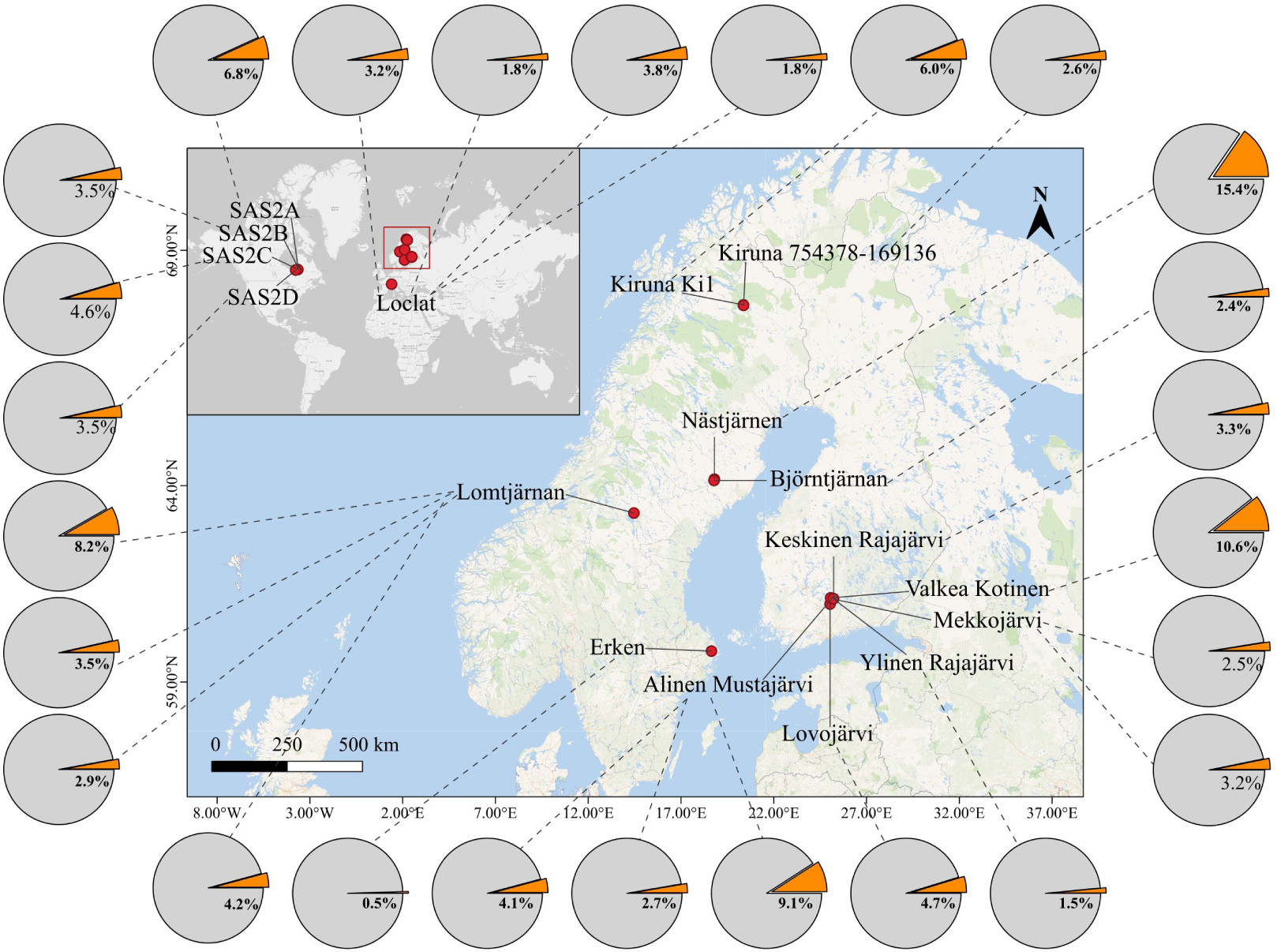
Distribution and relative abundance of MGC-containing bacteria. 267 samples were grouped into 80 lake-time groups by sampling sites and collection dates. 26 lake-time groups with more than 3 samples (N_t_ ≥ 3) and more than 2 MGC-containing samples (N_mtb_ ≥ 2) are selected for subsequent analyses. The ratio of the number of recruited reads to total metagenomic reads was used as a quantifier of MGC-containing community abundance in each sample. For each lake-time group, the sampled layer with the maximum MTB abundance value was considered the optimal niche for the corresponding MTB community. The map illustrates the geological distribution of the 26 lake-time groups. Pie charts around the map show the relative abundances of the MTB community in the 26 lake-time groups.

### Ubiquity and predominance of Deltaproteobacteria in freshwater MGC-containing communities

Since the discovery of MTB several decades ago, magnetotactic cocci within the Magnetococcia (also named as Etaproteobacteria) have long been considered as the most frequently observed and abundant MTB members in natural environments^29,30^. Contrary to this conventional view, here we found that MGC-containing members belonging to the Deltaproteobacteria were most ubiquitous in freshwater environments when samples were not pre-processed by magnetic enrichment (Fig. 5). Among the 165 samples from the filtered 26 lake-time groups, MGC-containing genomes were detected in 103 samples. Specifically, MGC-containing bacteria belonging to the Deltaproteobacteria, Alphaproteobacteria, and Magnetococcia were found in 76, 38, and 2 samples, respectively (Fig. 5a, Supplementary Fig. S2 and Supplementary Table S4). The distribution and relative abundance pattern of delta– and alphaproteobacterial MGC-containing bacteria within the 26 lake-time groups demonstrated the remarkable dominance of deltaproteobacterial MGC-containing bacteria (Fig. 5a). To determine whether the difference of relative abundances between MGC-containing Deltaproteobacteria and Alphaproteobacteria (with Magnetococcia excluded due to limited data) was significant, the nonparametric Mann-Whitney U test was employed (see Methods and Code Availability), which revealed that MGC-containing Deltaproteobacteria were significantly more abundant than MGC-containing Alphaproteobacteria within the 26 lake-time groups (*p*-value 0.0036). These results underlined the ubiquity and predominance of Deltaproteobacteria in freshwater MGC-containing communities.

**Fig. 5.**
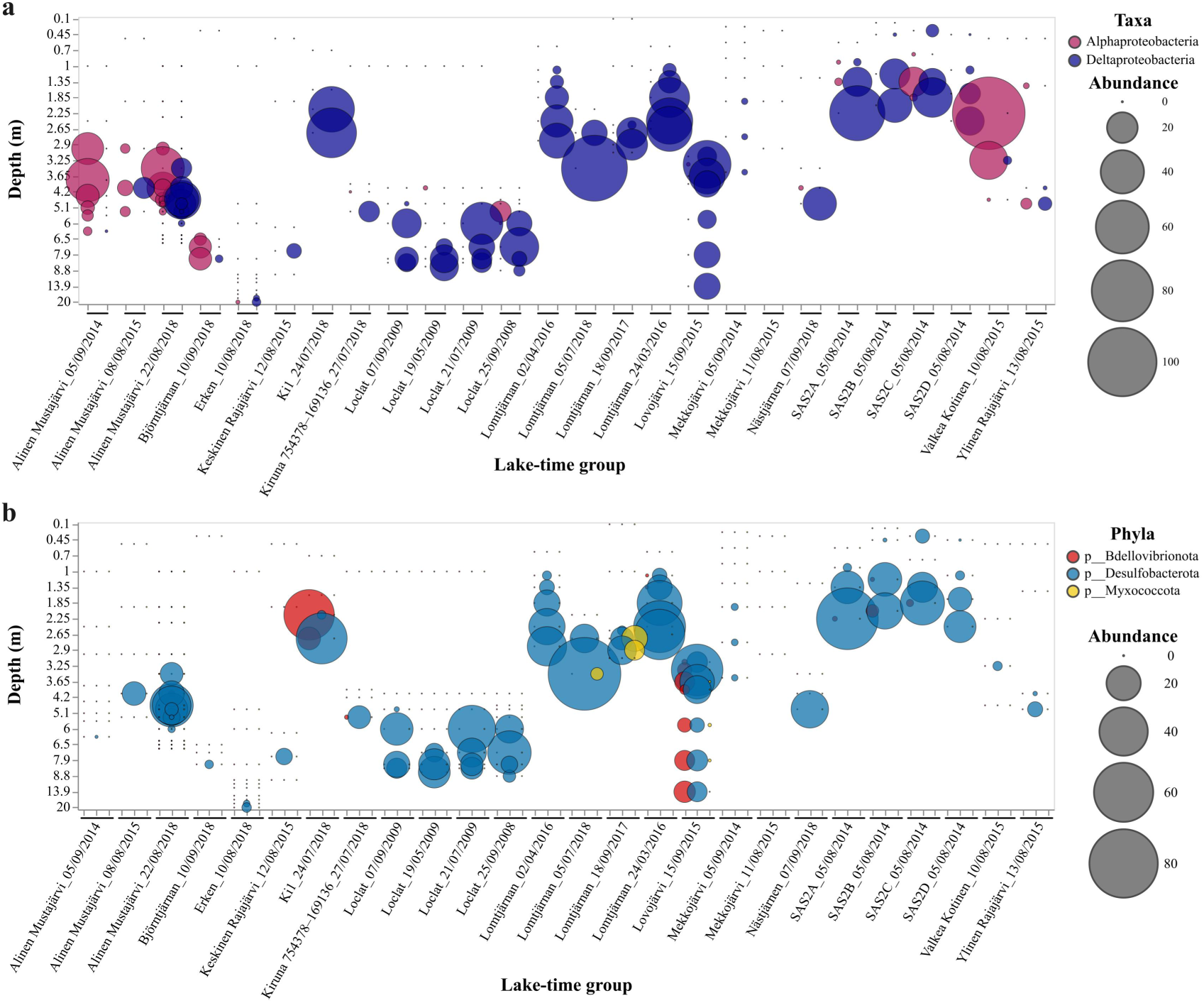
Distribution and abundance profile of MGC-containing Alphaproteobacteria and Deltaproteobacteria across 26 lake-time groups. Corrected coverage values of MGC-containing genomes within each metagenome were used as the abundance proxy (see methods for detail).For each metagenome, abundance values of MGC-containing genomes from the same lineage are summed. The X-axis represents 26 lake-time groups, and the Y-axis represents the sampling depth of each sample. Note that the Y-axis was arranged in ordinal order, but not scaled according to actual sampling depth. Bubble size represents the relative abundance of MGC-containing genomes affiliated with different lineages and the corrected coverage values were demonstrated on the right side of the bubble calibration. **a** Distribution and abundance profile of MGC-containing Alphaproteobacteria and Deltaproteobacteria across 26 lake-time groups. **b** Distribution and abundance profile of MGC-containing Deltaproteobacteria across 26 lake-time groups

To gain a better understanding of the MGC-containing Deltaproteobacteria within freshwater environments, we examined their distribution and abundance across 26 lake-time groups (Fig. 5b). Our analysis reveal a prevalence of Desulfobacterota within MGC-containing Deltaproteobacteria (note that phyla Desulfobacterota_G and Desulfobacterota_I are considered part of the phylum Desulfobacterota). MGC-containing Desulfobacterota were detected in 76 metagenome samples from 25 lake-time groups, followed by Bdellovibrionota that were identified in 16 samples from 7 lake-time groups and Myxococcota was observed in 6 metagenome samples from 3 lake-time groups (Fig. 5b). Recognized for their predatory lifestyle^39^, the non-negligible detection of MGC-containing Bdellovibrionota and Myxococcota genomes within the 26 lake-time groups investigated here highlights the potential role of magnetotaxis in bacterial predatory activities. In contrast, the absence of MGC-containing SAR32443 in this study suggests that SAR32443 may not be a common MTB population in freshwater environments. Collectively, these findings underscore the dominance of Desulfobacterota within the MGC-containing Deltaproteobacteria population in freshwater ecosystems.

### Lineage-specific vertical niche differentiation of MGC-containing bacteria in oxygen-stratified water columns

MTB are generally microaerophilic or anaerobic bacteria that inhabit near OATZs, where rapid decline in dissolved oxygen and significant changes in redox potential occur. Here, we profiled the vertical distribution of MGC-containing bacteria and analyzed their distribution patterns together with a group of environmental factors. To ensure a robust analysis, ten lake-time groups with sufficient physicochemical records (including dissolved oxygen, pH, electrical conductivity, temperature, ferric, ferrous, sulfate, nitrate, and ammonium) were selected for in-depth analysis (Fig. 6 and Supplementary Fig. S3). A distinct vertical distribution pattern of MGC-containing bacteria (gray bars in Fig. 6a and Supplementary Fig. S3) was observed across all lake-time groups, with the abundance peaking near or below the OATZs (shaded areas in Fig. 6a and Supplementary Fig. S3).

**Fig. 6.**
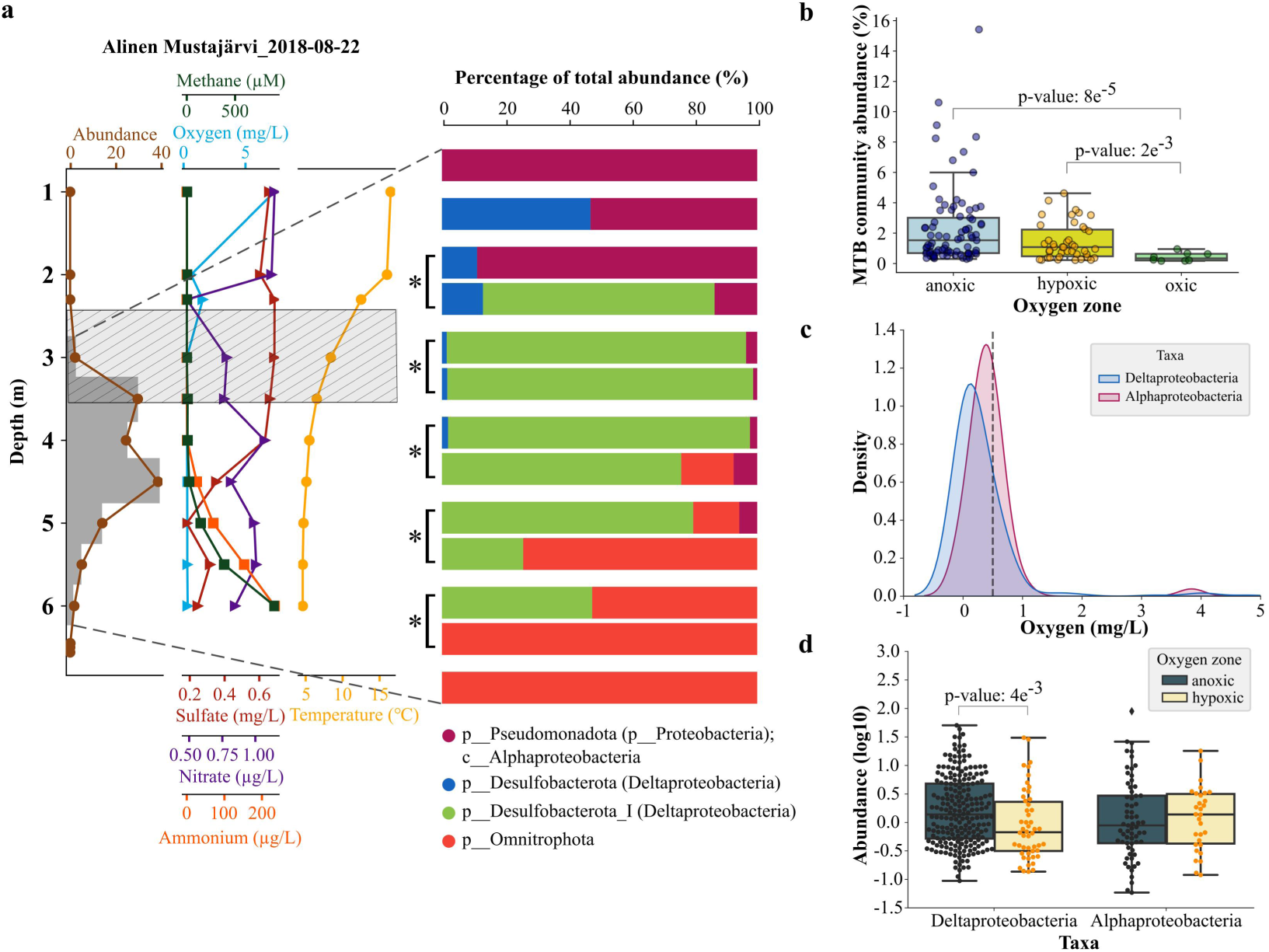
Vertical distribution of MGC-containing bacteria in a representative freshwater environment. **a** MGC-containing community profile along with the physicochemical factors of lake-time group Lake Alinen Mustajärvi_2018-08-22. Different physicochemical factors are illustrated with colored line charts. The grey bar chart on the left represents the relative abundance of MGC-containing community and the shaded area represents the oxic-anoxic transition zone (OATZ). MGC-containing bacterial taxonomy of each sample was shown on the right stacked bar chart, where different colors represent different taxonomic lineages. Samples with repeated sampling were illustrated with a left square bracket and a asterisk. **b** The relative abundances of MGC-containing community within samples from different niche zones (oxic, n=8; hypoxic, n=42; anoxic, n=81). Boxes represent the first quartile, median, and third quartile of abundance distribution values, and whiskers of 1.5 × interquartile range. Dots represent the relative abundance values in each individual sample.

The taxonomic compositions of MGC-containing bacteria were further investigated and distinct vertical shifts of dominant MGC-containing taxa were observed within the ten lake-time groups (Fig. 6a and Supplementary Fig. S3). For instance, MGC-containing Alphaproteobacteria dominated the upper layer of Lake Alinen Mustajärvi (Fig. 6a), while the dominant MGC-containing taxa shifted to Deltaproteobacteria and Omnitrophota in deeper layers, indicating a vertical niche-specific distribution of different MGC-containing taxa, possibly shaped by a combined effect of dissolved oxygen and different ion species.

To thoroughly characterize the vertical distribution of MGC-containing bacteria, the amount of dissolved oxygen was used to delineate the ecological niches of the collected samples, which were classified into three niche zones: samples with dissolved oxygen ≥ 2 mg/L were classified as the oxic zone, 0.5 mg/L < dissolved oxygen < 2 mg/L as the hypoxic zone (i.e., OATZ), and dissolved oxygen ≤ 0.5 mg/L as the anoxic zone. The relative abundance of MGC-containing bacteria in each niche zone was profiled (Fig. 6b), revealing that their relative abundance in hypoxic and anoxic zones are significantly higher than that in oxic zone. Although no significant difference was observed between hypoxic and anoxic zones, some anoxic metagenomes showed relatively high abundances of MGC-containing bacteria.

To further define the optimal ecological niches for major MGC-containing taxa, the distribution and relative abundance of 41 nr MGC-containing genomes were profiled in 131 MTB-containing samples which have dissolved oxygen records (Supplementary Fig. S4). The dominant MGC-containing Deltaproteobacteria and Alphaproteobacteria showed clear ecological niche differentiation. About three quarters of the MGC-containing Deltaproteobacteria occurred in the anoxic zones, whereas MGC-containing Alphaproteobacteria thrived in both hypoxic and anoxic samples (Fig. 6c and Supplementary Fig. S4). Moreover, the relative abundance of MGC-containing Deltaproteobacteria was significantly higher in the anoxic zone than in the hypoxic zone, while no significant difference was observed for MGC-containing Alphaproteobacteria between these two zones (Fig. 6d). These results indicate that MGC-containing Deltaproteobacteria are mainly thrived in the anoxic zone, whereas MGC-containing Alphaproteobacteria expanded from the hypoxic zone to the anoxic zone within the freshwater columns.

### Metabolic potential of dominant MGC-containing bacteria

Key metabolic pathways of the 41 nr MGC-containing genomes were reconstructed from their genomes (Fig. 7), and the metabolic potentials of the dominant MGC-containing members (Deltaproteobacteria and Alphaproteobacteria) within the freshwater environments investigated here were predicted and discussed below. The core carbohydrate metabolism pathways were present in all 41 nr MGC-containing genomes, of which five contained the complete carbon fixation pathways (Fig. 7). The Calvin–Benson–Bassham (CBB) pathway was found in MGC-containing Alphaproteobacteria and Gammaproteobacteria, coupled with the key CO_2_-fixing enzyme, ribulose-1,5-bisphosphate carboxylase/oxygenase (RuBisCO). Deltaproteobacterial MGC-containing bacteria could be furnished with other autotrophic metabolic strategies, such as the reductive tricarboxylic acid (rTCA) cycle pathway and the Wood-Ljungdahl (WL) pathway. Some MGC-containing Desulfobacterota genomes contain genes related to the WL pathway, which had previously been identified in Nitrospirota and Elusimicrobiota MTB genomes^40–42^, indicating that the WL pathway may be a common carbon fixation pathway in deep-branching lineages of autotrophic MTB.

**Fig. 7.**
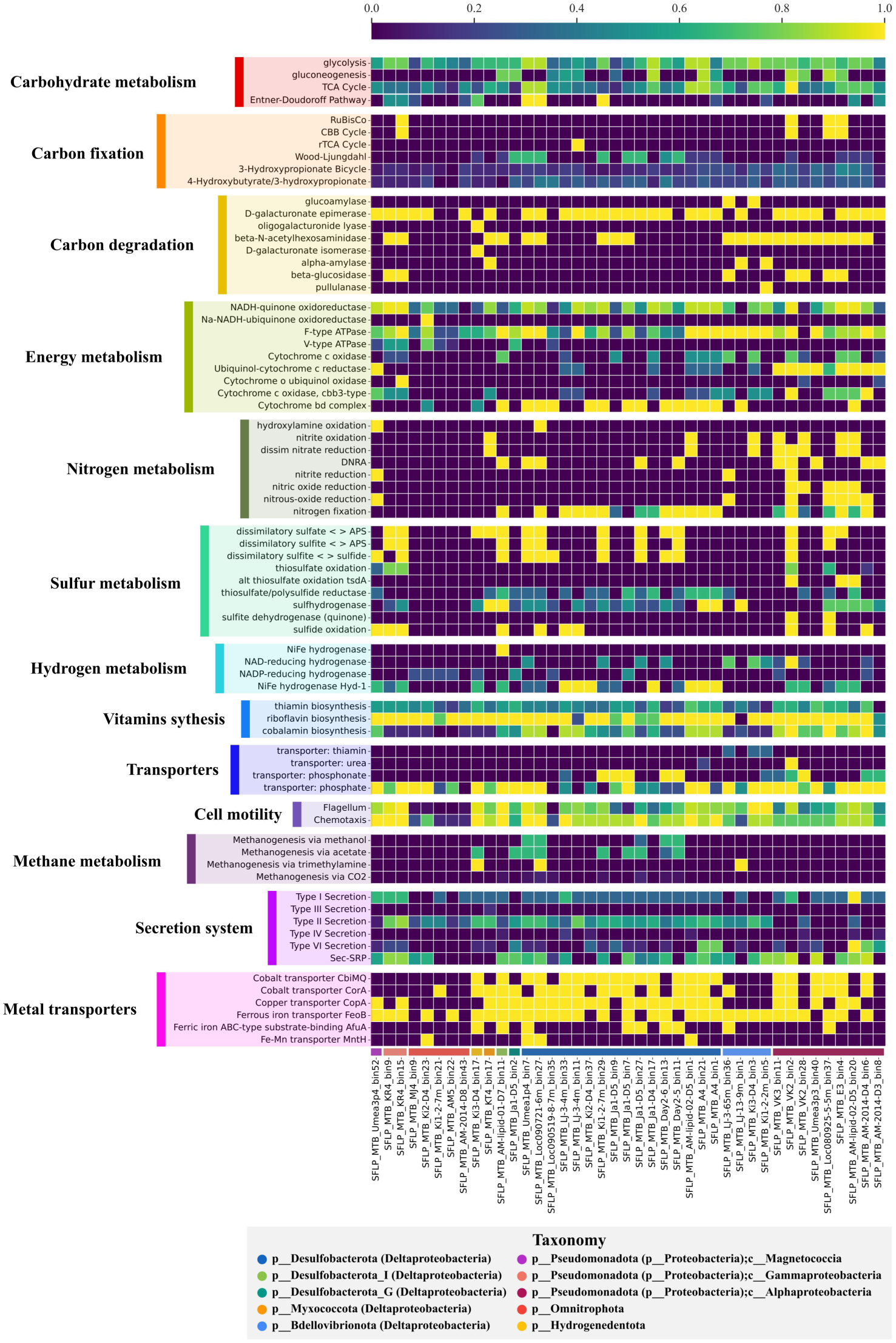
Metabolic prediction of 41 non-redundant MGC-containing genomes. The color gradient represents the completeness of major metabolic pathways inferred from the presence or absence of genes. Light represents a complete or nearly complete pathway, and dark represents a pathway that is absent or mainly incomplete.

Both MGC-containing Alphaproteobacteria and Deltaproteobacteria exhibit nitrogen fixation potential, as evidenced by the presence of the nitrogenase genes (*nifDKH*) within most of their genomes. Specifically, 14 out of 23 deltaproteobacterial MGC-containing genomes and 7 out of 9 alphaproteobacterial MGC-containing genomes contain *nifDKH* genes.

Key enzymes associated with the dissimilatory nitrate reduction to ammonium (DNRA) pathway, including nitrate reductases NarGHI and NapAB, were identified in both MGC-containing Alphaproteobacteria and Deltaproteobacteria, suggesting that both groups possess the capacity for anaerobic nitrate respiration via DNRA. However, the nitrite reductase enzymes differed between the two MGC-containing groups. NirBD was exclusively found in MGC-containing Alphaproteobacteria genomes, while NrfAH was exclusively found in MGC-containing Deltaproteobacteria genomes. Although both NirBD and NrfAH are dissimilatory nitrite reductases, NirBD was found to remain functional during aerobic growth in certain Actinomycetota^43^ and Gammaproteobacteria^44,45^ species. These findings imply that by empolying the NirBD enzyme, MGC-containing Alphaproteobacteria have the potential to conduct DNRA under aerobic conditions.

In addition to DNRA, genes associated with denitrification, including *nirK*/*nirS*, *norBC*, and *nosZ* (encoding nitrite, nitric oxide, and nitrous oxide reductases, respectively), were identified in both MGC-containing Alphaproteobacteria and Deltaproteobacteria genomes. This finding suggests that both groups possess the capability for complete denitrification under anaerobic conditions, using nitrate as the terminal electron acceptor.

The nitrite oxidoreductase (NxrAB), which is responsible for the final step in nitrification through catalyzing nitrite oxidation to nitrate^46^, was found in 4 out of 9 alphaproteobacterial MGC-containing genomes obtained in this study. Bacteria capable of nitrite oxidation are known as Nitrite-Oxidizing Bacteria (NOB). The genus *Nitrobacter* which is affiliated with Alphaproteobacteria is one of the well-known NOB populations^47^. However, the alphaproteobacterial nitrite oxidizers identified here belong to two families Magnetospirillaceae and Azospirillaceae (within the orders Rhodospirillales and Azospirillales, respectively), which are distinct from the genus Nitrobacter (which belongs to the family Xanthobacteraceae within the order Rhizobiales). Notably, the nitrite oxidation typically occurs in oxic environments suggests a dependency of MGC-containing Alphaproteobacteria on oxygen.

In terms of sulfur metabolism, a variety of genes encoding key enzymes related to assimilatory and dissimilatory sulfate reduction pathways were found in both MGC-containing Alphaproteobacteria and Deltaproteobacteria genomes, such as *Sat*, *CysND*, *CysC*, *CysH*, *CysJI*, *AprAB*, *Dsr*. However, we found that the thiosulfate metabolism within MGC-containing Alphaproteobacteria and Deltaproteobacteria are different, where *sox* genes encoding the sulfur oxidizing enzyme (SOX) system involved in the thiosulfate oxidation pathway exist in MGC-containing Alphaproteobacteria genomes and no *sox* genes were found within MGC-containing Deltaproteobacteria genomes. The Sox system from Alphaproteobacteria is well-established and studied^48^. Research on chemolithoautotrophic magnetospirillum magnetotacticum strains MV-1 and MV-2 revealed that they could grow microaerobically with S_2_O_3_^2-^ as an electron source and O_2_ as the electron acceptor^49^. Thus, the *sox* genes-bearing MGC-containing Alphaproteobacteria we found here might be able to harness energy from the oxidation of thiosulfate under aerobic conditions. On the other hand, MGC-containing Deltaproteobacteria are prone to perform thiosulfate reduction to sulfide as thiosulfate reductase (EC 1.8.5.5) was found to exist within 16 out of 23 MGC-containing Deltaproteobacteria genomes we obtained here. Thiosulfate reductase is typically found in anaerobic bacteria who perform anaerobic respiration and utilize thiosulfate as electron acceptors in the absence of oxygen^50^. Desulfovibrio species from the phylum Desulfobacterota are known sulfate-reducing bacteria that thrive in anaerobic environments^51^.

To summarize, we noticed that both MGC-containing Alphaproteobacteria and Deltaproteobacteria investigated here have the capability to anaerobic respiration, using nitrate or sulfate as electron acceptors. However, several metabolic traits that differ between these two groups revealed the possibility of MGC-containing Alphaproteobacteria for performing aerobic respiration in the presence of oxygen (hypoxic environment within the OATZ).

## Discussion

Due to their special abilities of magnetotaxis and prokaryotic organelle biosynthesis, MTB have long been of great interest. A deeper understanding of their distribution, abundance, and taxonomic composition would better unveil the ecophysiological importance of MTB communities in nature. It has long been considered that the overall abundance of MTB in microbiota is not very high (10^3^-10^6^ cells per mL)^16,17,30^ with some exceptions^20,52^, and for decades magnetotactic cocci within the class Magnetococcia (also known as Etaproteobacteria^29,53–55^) of the phylum Proteobacteria has been viewed as the most frequently identified and predominant MTB members in nature^17,29–31,56–62^. For example, as of January 2024, a total of 146 MTB or MGC-containing genomes affiliated to five bacterial phyla assembled from over 50 various freshwater environments have been reported, of which ∼42% (61 genomes) belong to the class Magnetococcia (Supplementary Table S5). Except for Magnetococcia, MTB belonging to the phylum Nitrospirota have occasionally identified as the predominant MTB groups in some lakes and acidic peatland soils^24,52^.

This study, however, challenges these conventional views and indicates that MGC-containing bacteria, that is, putative MTB, could represent unexpectedly high proportions (0.5% to 15.4% of total reads) in freshwater metagenomic data. Moreover, Deltaproteobacteria, rather than traditionally considered Magnetococcia and Nitrospirota, is the most ubiquitous and abundant MGC-containing group in freshwater columns. One possible reason for these findings could be that the abundance of whole MTB and the portion of Deltaproteobacteria in MTB communities have been underestimated in previous studies due to methodological bias. The typical pre-processing prior to most MTB studies was cell enrichment using a magnetic separation, such as capillary racetrack^63^ or MTB trap^64^. These magnetic enrichment strategies apply a magnetic field stronger than geomagnetic field to concentrate MTB cells from sediment or water samples relying on their magnetotaxis motility^33,65^. After enrichment, collected MTB cells are then utilized for subsequent analyses, such as morphological observations, 16S rRNA gene-based analysis, and omics-based analysis. During the magnetic enrichment step, fast-swimming cells with more efficient magnetotactic behavior, such as magnetotactic cocci within the Magnetococcia, are more preferentially enriched compared to those slow swimming populations with less efficient magnetotaxis^33^. Apart from magnetic cell enrichment, bias could also be introduced during the following 16S rRNA gene amplification or whole genomic DNA amplification processes^66–68^.

Alternatively, a recent study has revealed a silent biosynthetic MGC in a non-MTB alphaproteobacterial strain^69^. Unlike the majority of previous MTB studies, the 267 metagenomes in this study were directly obtained from water columns^34^, bypassing the magnetic enrichment steps. Thus, we cannot rule out the possibility that some of reconstructed MGC-containing genomes in this study may not represent active magnetosome-producing bacteria. Consequently, those bacteria with dormant MGCs could not be magnetically collected in conventional MTB studies. Although whether MGC-containing bacteria identified here are truly magnetotactic needs further investigations, considering that transcriptionally silent magnetosome genes may have the potential to be activated under certain conditions^69^, the unexpectedly high relative abundance of MGC-containing bacteria in freshwater microbiota and the predominance of Deltaproteobacteria in MGC-containing communities not only improve our understanding of diversity and ecology of putative MTB but also highlight the previous underestimation of their roles in global biogeochemical cycling in oxygen-stratified habitats. Moreover, newly identified MGCs from different taxonomic lineages in this study provide valuable functional gene resources for future production of transgenic biogenic magnetic nanoparticles with various sizes, shapes, and physicochemical properties in foreign hosts^70–72^.

To determine the mechanisms and environmental conditions supporting MGC-containing bacteria in the oxygen-stratified water column, genome-wide quantitative read recruitment has been used to decipher their vertical distribution and microbial function across gradients of redox substrates. This study reveals a stratification of MGC-containing bacteria in water columns followed a niche differentiation pattern. Taking advantage of genome-based metabolic prediction, our results delineate clear associations between metabolic strategies and ecological niches of MGC-containing bacteria, especially the two dominant MGC-containing groups of Deltaproteobacteria and Alphaproteobacteria. The relative abundances of MGC-containing Alphaproteobacteria and Deltaproteobacteria are significantly correlated with several physicochemical parameters (Supplementary Fig. S5), which should play important roles in shaping their vertical distribution. MGC-containing Alphaproteobacteria live in hypoxic to anoxic water columns that have the capacity for thiosulfate oxidation and nitrite oxidation under the presence of oxygen. While, the occurrence of MGC-containing Deltaproteobacteria which conduct anaerobic respiration is mainly locate in the anoxic layers. Considering their high abundance in anoxic water columns, our results indicate that MGC-containing Deltaproteobacteria should play a greater role in the carbon, sulfur, nitrogen, and iron biogeochemical cycles in oxygen-deficient aquatic regions than previously thought.

One striking finding in this study is that we do not identify any Fe_3_S_4_-type MGC-containing genomes near or below the OATZ, which is in contrary to the classical vertical distribution model of MTB that Fe_3_S_4_-producing populations normally occur in anoxic conditions^20,73,74^. It is likely that the abundance of Fe_3_S_4_-producing MTB in freshwater lakes and ponds analyzed here is too low to be recovered using genome-resolved metagenomics applied in this study. Therefore, magnetic enrichment pre-processing may be more appropriate for investigations of low-abundance Fe_3_S_4_-producing MTB in nature.

In recent years, a growing number of MTB-containing phyla have been discovered, suggesting that our knowledge on MTB taxonomy remains limited^24,25,41,54^. In the present study, a total of 63 MGC-containing genomes have been reconstructed, expanding the genomic and taxonomic diversity of putative MTB. These genomes belong to eight bacterial phyla, including the first genomic evidence for the MGC-containing population within the phylum Myxococcota. Bacteria in Myxococcota are well known for their facultative predatory lifestyle^75^. In addition to Myxococcota, four MGC-containing genomes from the obligate predators Bdellovibrionota^76^ are obtained here. Combined with a previous study^24^ that reconstructed six Bdellovibrionota MTB genomes, these findings suggest that MGCs and thus putatively magnetotaxis might be widely distributed among predatory bacteria. With broad prey spectra, predatory Myxococcota and Bdellovibrionota are hypothesized to play currently unknown roles in the microbial food web^77^, and bacterial predation may serve as a previously unrecognized selective force in microbial ecosystems^39^. MTB of a predatory lifestyle may take advantage of the geomagnetic field to efficiently prey on bacteria dwelling in or near the OATZ regions and further influence the bacterial communities in their respective habitats. Further investigation into potentially predatory MTB will help understand their ecological roles in regulating microbial community structures, prey populations, and nutrient cycling.

In summary, our results reveal an unexpectedly high relative abundance of MGC-containing bacteria in freshwater microbiota and the ubiquity and predominance of Deltaproteobacteria in MGC-containing communities in freshwater ecosystems. This study emphasizes that our understanding of abundance, diversity and distribution of putative MTB in nature is still limited even after several decades of research, thus their ecological roles in aquatic biogeochemical cycling remain to be evaluated.

## Methods

### Shotgun metagenomes

A total of 267 shotgun metagenomes under the project accession PRJEB38681 were downloaded from the European Nucleotide Archive (ENA). These metagenomes were sampled and reported in Buck et al^34^ (see Supplementary Table S1). All metagenomes were sequenced on the Illumina NovaSeq platform using a paired-end 2 × 150 bp sequencing strategy at the Science for Life Laboratory (Uppsala University, Uppsala, Sweden)^34^.

### Reads quality control, metagenomic assembly, binning, and bin refinement

Raw reads of all metagenomes were trimmed and filtered using the Read_qc module of metaWRAP v1.1.5^78^ with the ‘--skip-bmtagger’ flag. For each metagenome, clean reads were then assembled using metaSPAdes v3.13.0^79^ with the minimum length of assembled contigs set to 2000 bp (‘-l 2000’). Assemblies were then binned with the metaWRAP Binning module using three binnners: metaBAT2 v2.12.1^80^, MaxBin2 v2.2.6^81^, and CONCOCT v1.0.0^82^. For each metagenome, the three original bin sets were then compared, and a refined bin set was obtained using the metaWRAP Bin_refinement module with the minimum completion of bins set to 50% and the maximum contamination set to 10% (‘-c 50 –x 10’).

### Annotation and identification of MGC-containing genomes

We employed MagCluster v0.2.2^83^ to annotate the refined bins and screen for putative magnetosome gene clusters (MGCs). Briefly, (i) bins were first annotated by MagCluster using Prokka v1.13.3^84^ with the e-value set to 1e^−05^; (ii) mgc_screen module of MagCluster was then used to screen for putative MGCs with default parameters; (iii) putative magnetosome protein sequences were then retrieved and compared against NCBI non-redundant protein sequence database using NCBI Blast+ suite v2.12.0 (ftp://ftp.ncbi.nlm.nih.gov/blast/executables/blast<u>+/</u>) using the Position-Specific Iterated (PSI-BLAST) algorithm with the number of iterations set to 1 (-num_iterations 1); (iv) The PSI-BLAST results of all putative magnetosome proteins were manually checked; and (v) MGCs were visualized using clinker v0.0.23^85^.

### Quality assessment of MGC-containing genomes

The quality (completeness and contamination) of MGC-containing genomes was estimated with CheckM v1.0.12^86^ using the ‘lineage_wf’ workflow. Genomic statistics, including genome size, GC content, contigs N50/L50, and maximum contig length, were analyzed using QUAST v5.0.2^87^.

### Dereplication of redundant MGC-containing genomes

A non-redundant MGC-containing genomic dataset was obtained using dRep v3.4.0^88^. Note that the genomes’ quality filter was set to a minimum completeness of 50% and maximum contamination of 10% (the medium-quality MAGs based on the MIMAG standard^35^). ANImf algorithm was used for ANI computation. The ANI thresholds for primary clusters and secondary clusters were set to 0.9 and 0.99, respectively. After dereplication, representative MGC-containing genomes generated a non-redundant dataset for further analysis.

### Taxonomic classification and phylogenomic analysis of MGC-containing genomes

Taxonomic classification of 41 non-redundant MGC-containing genomes was performed using the Genome Taxonomy Database Toolkit (GTDB-Tk) v2.1.0^89^ with database release 207_v2 using the ‘classify_wf’ workflow. Phylogeny of these genomes was analyzed together with 33 previously published MTB genomes and 60 non-MTB genomes. For each genome, the 120 bacterial single-copy marker protein^90^ sequences were identified, aligned, and concatenated using GTDB-Tk. The concatenated sequences were then used to infer the phylogeny. The phylogenomic tree was constructed using FastTree v2.1.10^91^ under the WAG model (‘-wag’). The phylogenomic tree was rooted with three genomes of the phylum Patescibacteria (NCBI accession number MFYJ01000001, MNWP00000000, CAIKAD010000001) using Interactive Tree Of Life (iTOL) v6^92^. The tree was then visualized together with GC content, genome size, completeness, and contamination data using anvi’o v7.1^93^.

### Profiling of MGC-containing genomes across all metagenomes

The 41 non-redundant MGC-containing genomes were used for distribution and abundance profiling of putative MTB in all metagenomes by conducting a read recruitment procedure using anvi’o v7.1 as described in Delmont et al^36^. Briefly, all contigs and scaffolds from the non-redundant MGC-containing genomes were first concatenated into a single FASTA file. Bowtie2 v2.3.5^94^ was next used to build the index files. Short reads from each metagenome were then mapped against all contigs and scaffolds with the ‘--no-unal’ parameter, and the alignment results were stored as SAM files. Then, samtools^95^ was used to convert the SAM files to BAM files. anvi’o v7.1 was used to profile each BAM file to estimate the coverage and detection statistics of each scaffold, followed by combining all mapping profiles into a merged profile database for each metagenome. Briefly, the ‘anvi-profile’ program was used to calculate the coverage value per nucleotide position of each BAM file and stored as an anvi’o profile database. The ‘anvi-merge’ program was used to convert all profile databases into a merged profile database, and the information of contigs or scaffolds was imported into their corresponding MTB MAGs of the profile database using the ‘anvi-import-collection’ command. Then ‘anvi-summarize’ was used to report the statistics of read recruitment results, including the coverage and detection data for each MGC-containing genome across all metagenomes.

### Characterization of the presence and abundance of putative MTB across all metagenomes

The detection and coverage statistics of non-redundant MGC-containing genomes across all metagenomes were calculated after read recruitments by anvi’o v7.1 (Supplementary Table S3). The detection value was used to demonstrate whether MGC-containing genomes existed in metagenomes, and the presence of genomes was confirmed when detection ≥ 0.25^38^. To eliminate the by-effects of non-specific mapping during read recruitments, nucleotide positions whose coverage values fall within the 1st and 4nd quartiles were not considered, and the interquartile mean coverage value was used as a proxy of the relative abundance of putative MTB. When the detection value of a MGC-containing genome was smaller than 0.25, the interquartile mean coverage value was set to zero.

### Metabolic characterization prediction

Gene function annotation was first conducted using KofamScan v1.3.0^96^ by mapping protein sequences against the HMMs within the KOfam database, and KO identifiers were assigned to each gene. By default, an asterisk ‘*’ is added if the mapping score is higher than the predefined threshold by KofamScan (see https://github.com/takaram/kofam_scan). Significantly mapped genes against the KOfam database were filtered out using a bash script (kofam.get.sig.sh, see https://github.com/RunJiaJi/MTBProfiling/). The completeness of key metabolic pathways was estimated and illustrated by KEGGDecoder v1.3^97^. Pathway maps were then generated using the online KEGG Mapper Reconstruct tool (https://www.genome.jp/kegg/mapper/reconstruct.html) and manually checked.

### Data processing and visualization

All data processing and statistical analyses were performed in the interactive development environment of JupyterLab v3.3.2 (https://jupyter.org/). Python v3.9.5 (https://www.python.org/) with several libraries and packages were used. Subsequently, pandas v1.3.4 (https://pandas.pydata.org/) and numpy v1.22.4 (https://numpy.org/) were used for data organization, integration, and cleaning. Spearman Correlation analysis between MTB abundance and environmental factors was conducted using scipy v1.8.1 (https://scipy.org/) with the scipy.stats.spearmanr function. The map of sample sites was generated in QGIS v3.30 (https://qgis.org/en/site/). Data visualization for figures was conducted using matplotlib v3.5.2 (https://matplotlib.org/), altair v4.2.0 (https://altair-viz.github.io/), and seaborn v0.11.2 (https://seaborn.pydata.org/). All figures were finally organized using the open-source vector graphics editor Inkscape v1.2.2 (https://inkscape.org/).

## Data availability

The raw metagenomic sequencing data are available in the European Nucleotide Archive (ENA) under project PRJEB38681. All MGC-containing genomes reconstructed in this study have been deposited in the NCBI BioProject under PRJNA400260 (accession numbers pending).

## Code availability

Codes and scripts used in the metagenomic analysis and data processing are freely available under the MIT License at https://github.com/RunJiaJi/MTBProfiling

## Acknowledgements

We gratefully acknowledge Moritz Buck for permission to use the metagenomic data they reported. We thank the Supercomputing Laboratory of the Institute of Geology and Geophysics, Chinese Academy of Sciences, for their invaluable support in data transfer and storage. We thank Wensi Zhang for his help on metagenomic dataset download. We thank Jia Liu for her advice on drafting figures. We thank Yinzhao Wang on his advice on the metabolism analysis. This work was supported by the National Natural Science Foundation of China (NSFC) grants 42388101, 42293293 and T2225011.

## Author contributions

W.L. and R.J.J. designed the study. R.J.J. conducted the metagenomic analysis and statistical analysis. R.J.J. and W.L. drafted the manuscript with input from J.X.S., J.W., and Y.X.P. All authors contributed to critical revisions and approved the final manuscript.

## Additional information

**Supplementary information** The online version contains supplementary material.

