## Supplementary figures and images for "The under-recognized dominance of magnetosome gene cluster-containing bacteria in oxygen-stratified freshwater ecosystems"

### Supplementary Fig. S1

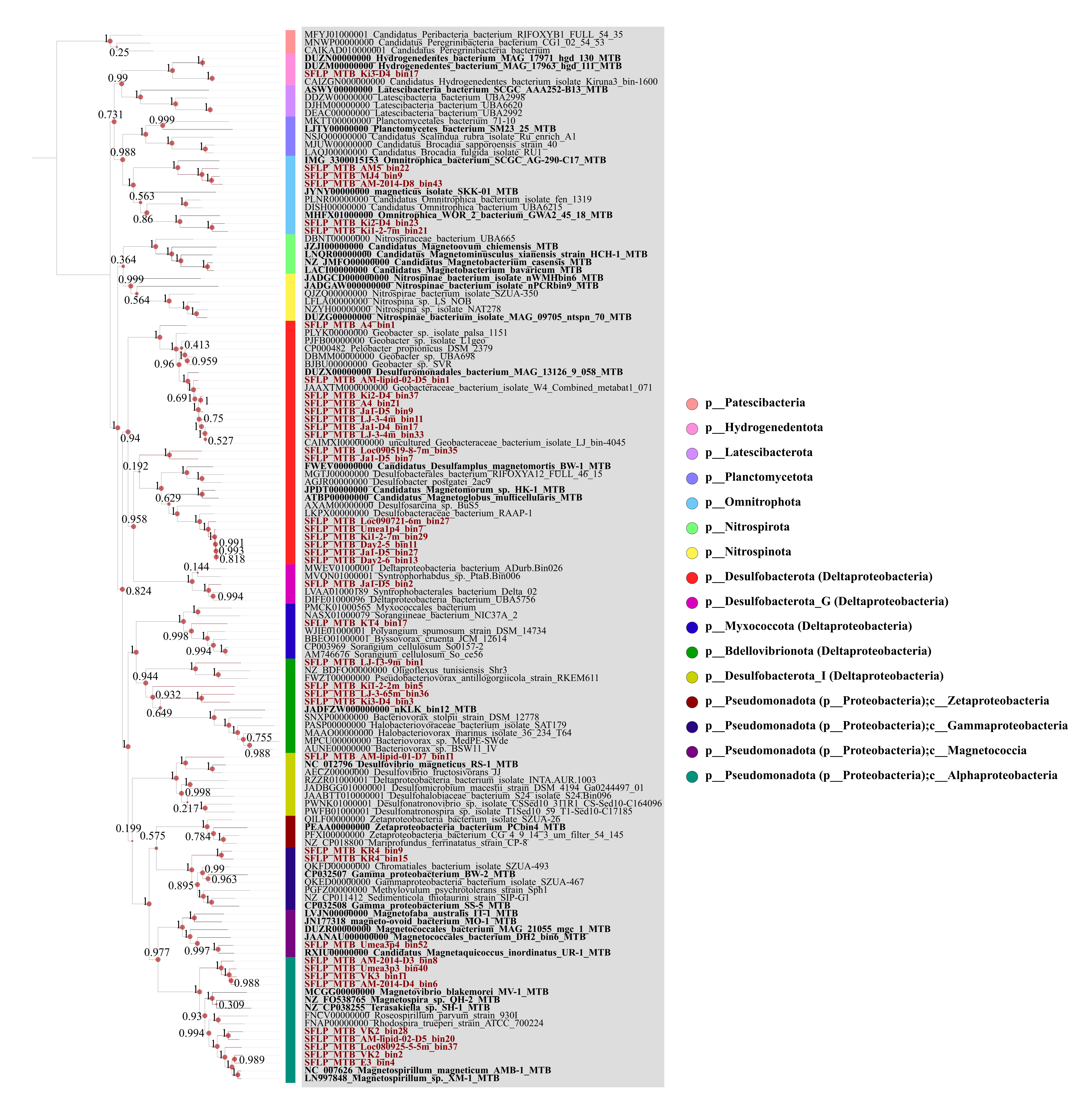

### Supplementary Fig. S2

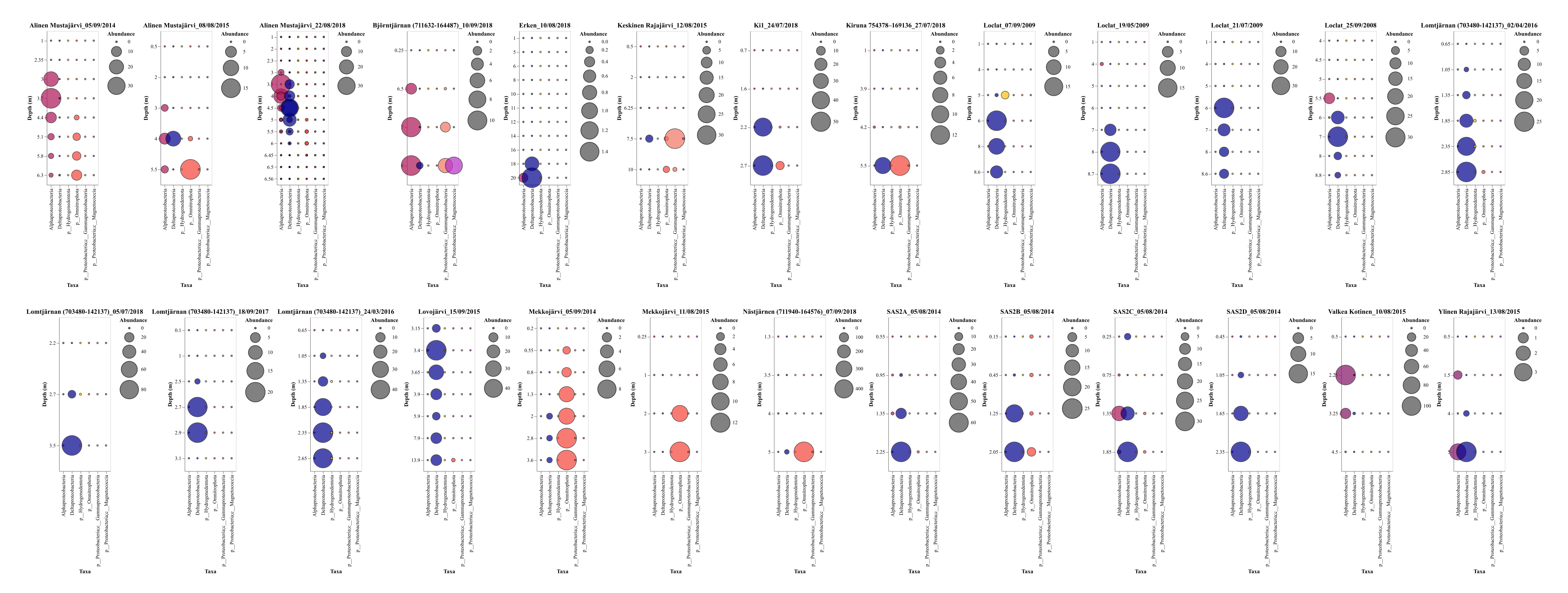

### Supplementary Fig. S3

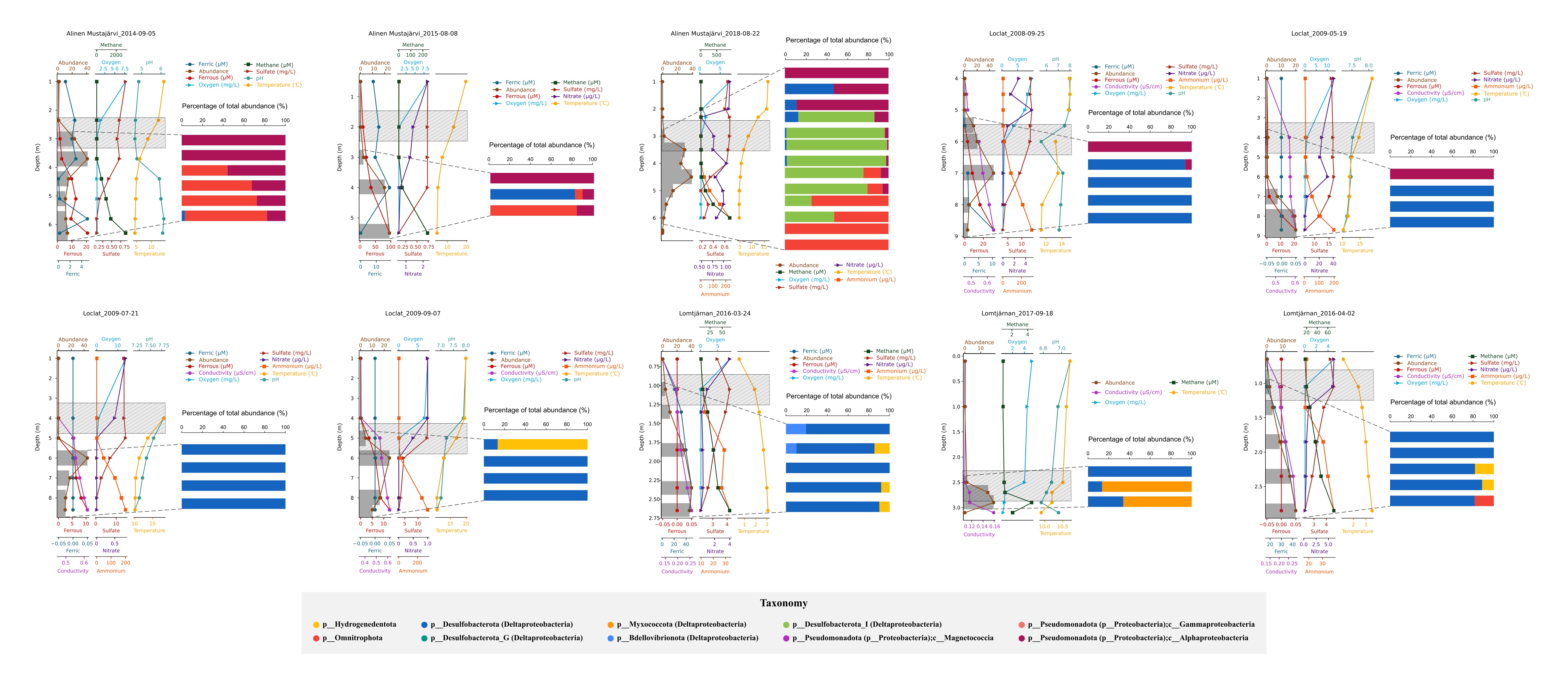

### Supplementary Fig. S4

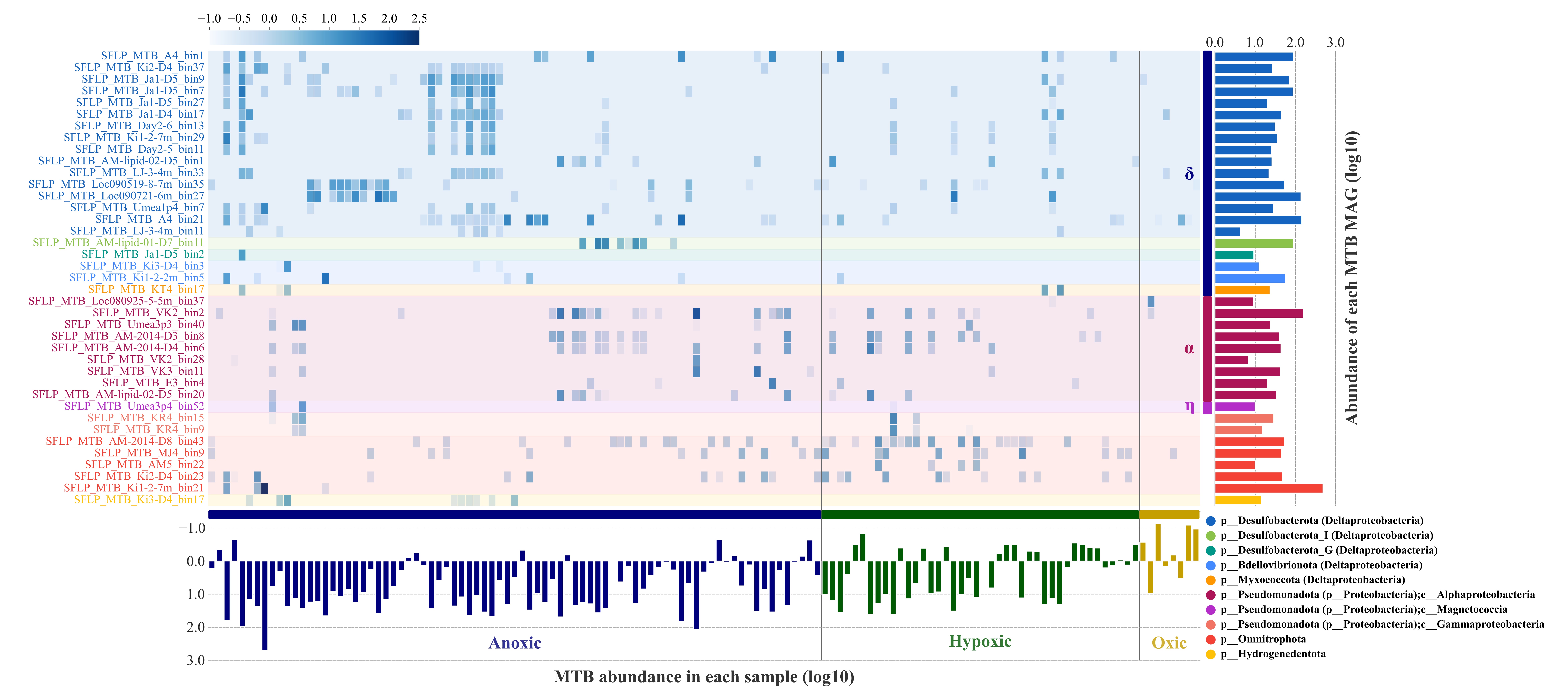

### Supplementary Fig. S5

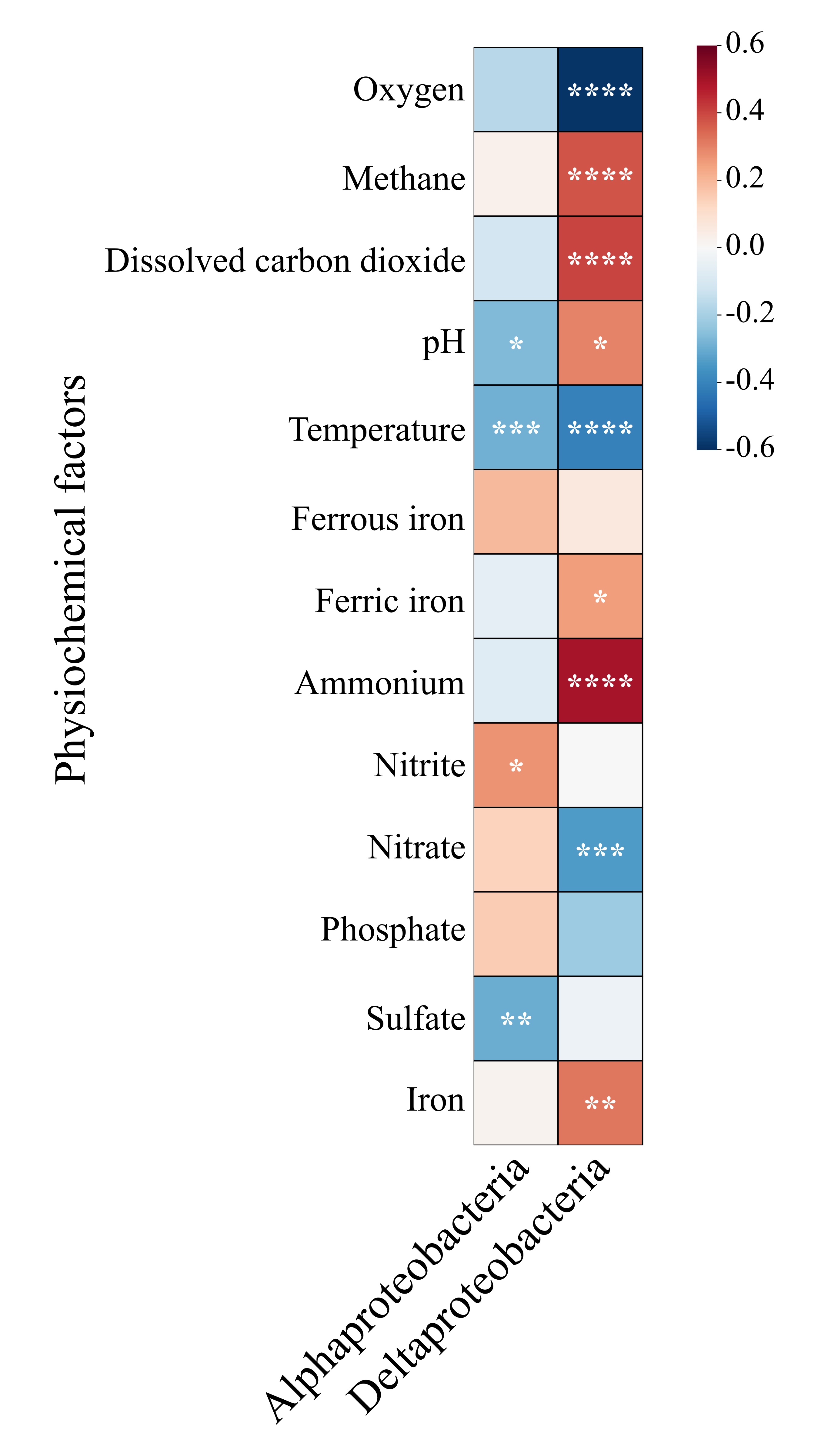
